# Acute Exercise Following Skill Practice Promotes Motor Memory Consolidation in Parkinson’s Disease

**DOI:** 10.1101/2020.05.15.097394

**Authors:** Philipp Wanner, Martin Winterholler, Heiko Gaßner, Jürgen Winkler, Jochen Klucken, Klaus Pfeifer, Simon Steib

## Abstract

Acute cardiovascular exercise has shown to promote neuroplastic processes, and thus to improve the consolidation of newly acquired motor skills in healthy adults. First results suggest that this concept may be transferred to populations with motor and cognitive dysfunctions. In this context, Parkinson’s disease (PD) is highly relevant since patients demonstrate deficits in motor learning. Hence, in the present study we sought to explore the effect of a single post-practice exercise bout on motor memory consolidation in PD patients.

For this purpose, 17 PD patients (Hoehn and Yahr: 1 – 2.5, age: 60.1 ± 7.9 y) practiced a whole-body task followed by either (i) a moderate-intense bout of cycling, or (ii) seated rest for a total of 30 minutes. The motor task required the participants to balance on a tiltable platform (stabilometer) for 30 seconds. During skill practice, patients performed 15 trials followed by a retention test 1 day and 7 days later. We calculated time in balance (platform within ± 5° from horizontal) for each trial and within- and between-group differences in memory consolidation (i.e. offline learning = skill change from last acquisition block to retention tests) were analyzed.

Groups revealed similar improvements during skill practice (F_4,60_ = .316, p = .866), but showed differences in offline learning, which was only evident after 7 days (F_1,14_ = 5.602, p = .033).

Our results suggest that a single post-practice exercise bout is effective in enhancing long-term motor memory consolidation in a population with motor learning impairments. This may point at unique promoting effects of exercise on dopamine neurotransmission involved in memory formation. Future studies should investigate the potential role of exercise-induced effects on the dopaminergic system.

**Highlights:**

- Acute exercise enhanced motor memory consolidation in PD
- Effects were evident only at 7-day retention
- Results may indicate unique exercise-effects on the dopaminergic system
- Findings show promising potential of exercise for motor rehabilitation

## 1 Introduction

There is no doubt that cardiovascular exercise promotes brain health and function (Cotman, 2002; Dishman et al., 2006; Hillman, Erickson, & Kramer, 2008). Remarkably, already a single bout of exercise has shown to facilitate the stabilization (i.e. memory consolidation) of newly acquired motor skills (Roig et al., 2016; Taubert, Villringer, & Lehmann, 2015). This process of motor skill learning relies on functional and structural brain plasticity (Dayan & Cohen, 2011; Doyon et al., 2009). Accordingly, first mechanistic models linked the behavioral findings to exercise-induced effects on several neuroplastic processes (Basso & Suzuki, 2017; El-Sayes, Harasym, Turco, Locke, & Nelson, 2018; Moriarty et al., 2019; Roig et al., 2016; Taubert et al., 2015). Briefly, this comprises the upregulation of catecholamines (e.g., dopamine) as well as neurotrophic factors (e.g. BDNF) (Dinoff, Herrmann, Swardfager, & Lanctôt, 2017; McMorris, Collard, Corbett, Dicks, & Swain, 2008), positive effects on the primary motor cortex as well as synaptic plasticity (Mellow, Goldsworthy, Coussens, & Smith, 2019; Singh & Staines, 2015), and an increased cerebral metabolism (Ogoh & Ainslie, 2009).

While current experiments mainly focused on healthy young adults, the effects of acute exercise may also be relevant for rehabilitation purposes (Charalambous et al., 2018; Nepveu et al., 2017). In this context, exercise might be a promising complementary non-pharmacological treatment for individuals with Parkinson’s disease (PD). PD is a neurodegenerative disorder characterized by a loss of dopaminergic neurons in the caudal basal ganglia. In addition to motor symptoms (i.e. bradykinesia, tremor, rigidity, and postural instability), the depletion of dopamine results in cognitive deficits, including impaired memory formation (Moustafa et al., 2016). A key component of PD motor rehabilitation is the repeated practice of motor skills (e.g. balance, gait), and therefore can be considered as motor (re)learning processes (Abbruzzese, Marchese, Avanzino, & Pelosin, 2016). However, the basal ganglia (i.e. corticostriatal circuits) are involved in motor learning, particularly during memory consolidation (Doyon et al., 2009). Consequently, PD patients demonstrate diminished skill retention compared to nondisabled individuals of similar age (Marinelli, Quartarone, Hallett, Frazzitta, & Ghilardi, 2017; Nieuwboer, Rochester, Müncks, & Swinnen, 2009; Olson, Lockhart, & Lieberman, 2019). While pharmacological treatment appears not to enhance motor learning adequately (Marinelli et al., 2017), exercise may serve as an intervention to counteract these deficits. Remarkably, Petzinger and colleagues (2013) recently proposed a model for motor skill practice (i.e. motor learning) and exercise working synergistically to optimize neuroplastic processes in PD. Furthermore, due the dopamine depletion, PD is an ideal model to gain deeper insight into the effects of exercise on the dopaminergic system.

The synergistically working mechanisms of skill practice and exercise may explain the sustainable effects of treadmill training with postural pertubations in PD as seen in our experiments (Steib et al., 2017, 2019). In accordance to the practice-exercise interaction, treadmill walking combines intense skill practice with an exercise stimulus, thereby facilitating the consolidation of the motor skill. In line with this, we recently observed that moderate intense cycling preceding skill practice improves offline learning compared to seated rest in PD patients (Steib et al., 2018). While online gains during memory encoding were not enhanced, groups showed differences in the time course of skill improvements during practice. Specifically, exercise resulted in larger initial, but reduced late online gains, which might be the consequence of fatigue (Roig, Skriver, Lundbye-Jensen, Kiens, & Nielsen, 2012). Our results, therefore, provide first evidence for promoting effects of acute exercise on motor memory consolidation in PD. However, additional research is necessary to confirm these findings and explore whether the improved offline gains are only an artifact of fatigue during the end of skill practice (Robertson, 2019).

Interestingly, first work in healthy populations demonstrated improved motor memory consolidation with exercise not only preceding, but also performed following skill practice (Dal Maso, Desormeau, Boudrias, & Roig, 2018; Ferrer-Uris, Busquets, Lopez-Alonso, Fernandez-Del-Olmo, & Angulo-Barroso, 2017; Lundbye-Jensen, Skriver, Nielsen, & Roig, 2017; Ostadan et al., 2016; Thomas, Flindtgaard et al., 2016; Thomas, Johnsen et al., 2016) and these effects were even stronger compared to pre-practice regimes (Roig et al., 2012; Tomporowski & Pendleton, 2018). Consequently, it is proposed that exercise preceding skill practice mainly facilitates the phase of memory encoding and early stages of consolidation, whereas post-practice exercise exclusively primes the phase of memory consolidation (Roig et al., 2016).

Taken together, skill practice (i.e. motor learning) is a core component of non-pharmacological treatment in PD, but motor memory consolidation is impaired in this population. First results emphasize acute exercise as a promising intervention to counteract these deficits, but pre-practice exercise may cause fatigue during memory encoding. Moreover, the dopamine denervation in PD is an ideal model to test a potential dopaminergic stimulation by exercise, and thereby providing a better understanding of the underlying mechanisms. Thus, in the present pilot study we aimed at investigating the effects of post-practice exercise on motor memory consolidation in PD patients. We hypothesized that moderate intense cycling immediately following skill practice improves offline learning compared to seated rest.

## 2 Methods

This study was preregistered (registration number: NCT03886090) at ClinicalTrials.gov. The registration protocol is accessible at https://clinicaltrials.gov/ct2/show/NCT03886090.

### 2.1 Participants

A sample of 18 early to mid-stage PD patients participated in this study. Patients were recruited at two different hospitals where they underwent medical treatment. We included patients if they met the following criteria i) a Hoehn & Yahr score of < 3; ii) a score of ≤ 1 in the Unified Parkinson’s Disease Rating Scale (UPDRS) item ‘postural stability’; iii) were able to stand and walk independently; and iv) were unfamiliar to the motor task. Criteria for exclusion were i) higher level of cognitive impairment indicated by a score of < 21 in the Montreal Cognitive Assessment (MoCA) (Dalrymple-Alford et al., 2010); ii) other clinically relevant neurological, internal or orthopedic conditions besides Parkinsonism that would interfere with the exercise paradigm or motor learning task; iii) musculoskeletal conditions or surgery three months before the study enrolment.

All experiments were conducted in accordance with the Declaration of Helsinki, the study was approved by the local ethics committee (reference number: 125_17B), and patients gave written informed consent prior to participation.

### 2.2 Experimental design

Participants attended a total of four sessions inculding i) a pre-examination not more than 14 days prior to; ii) an acquisition session where the motor task was practiced, followed by iii) a retention test 24 ± 2 hours, and iv) 7 days ± 2 hours later (figure 1). During the acquisition session, the participants were allocated after block randomization into one of two groups performing either i) continuous moderate-intensity exercise (EX), or ii) seated rest (REST) for a total of 30 minutes immediately (≤ 5 minutes) after practicing the motor task. The block randomization was stratified by gender (male / female) and age (< 62 / ≥ 62), since these two factors may modulate the effects of exercise on cognitive performance (Smith et al., 2005) and memory formation (Kamijo et al., 2009; Kramer & Colcombe, 2018). Participants were blinded to the researchers’ hypotheses regarding the effect of the different conditions. Furthermore, PD patients were instructed to refrain from vigorous physical activity 24 hours prior to the acquisition as well as retention sessions.

**Figure 1.**
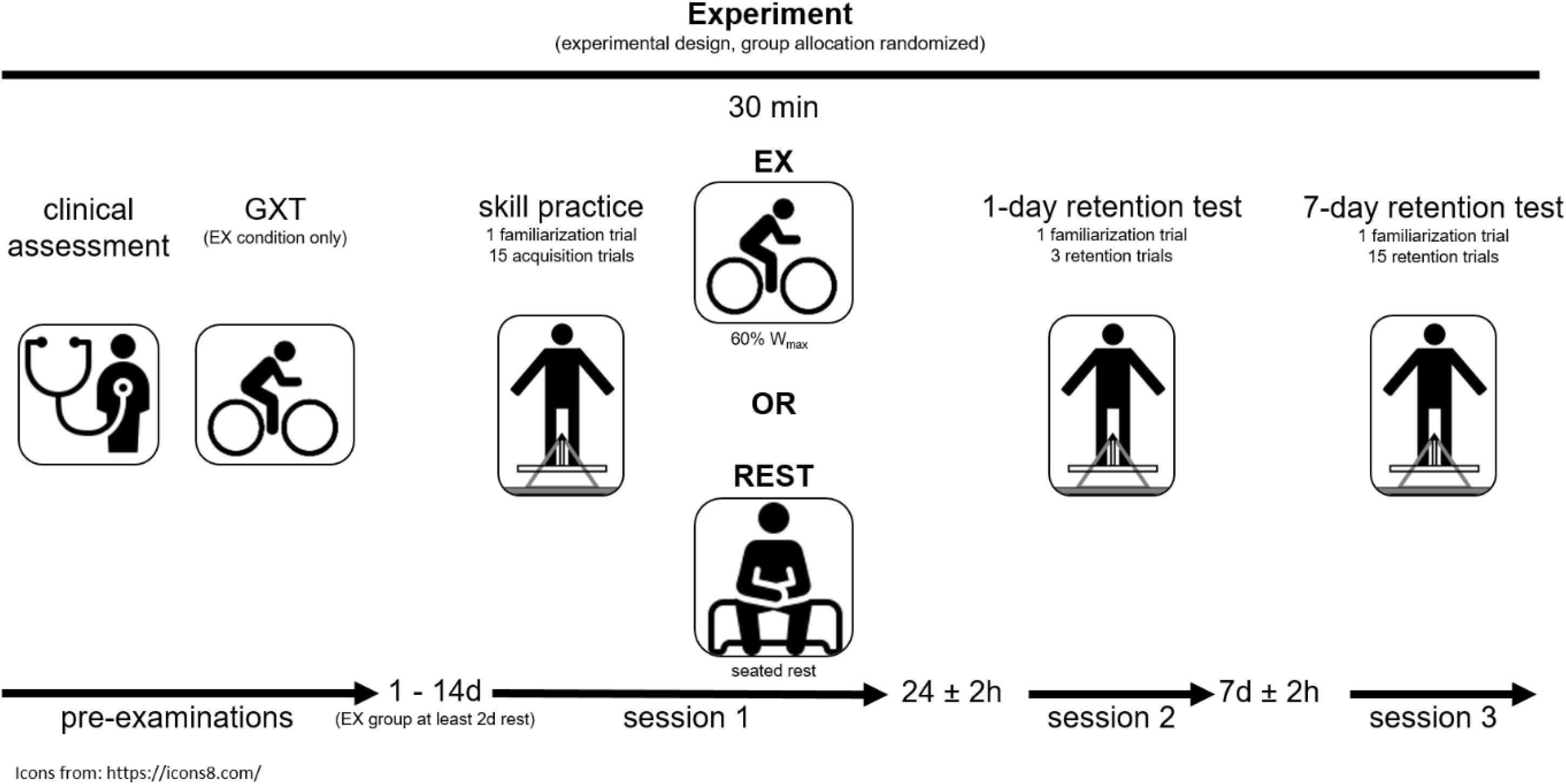
Schematic overview of the experimental design; GXT = graded exercise test; EX = exercise.

### 2.3 Pre-examination

The pre-examination was carried out not more than 14 days prior to the main experiment. The session included the assessment of anthropometric and demographic data, motor function using the motor part of the Unified Parkinson’s Disease Rating Scale (MDS-UPDRS-III; Goetz et al., 2008), cognitive function using the MoCA (Dalrymple-Alford et al., 2010), and self-reported physical activity level using the International Physical Activity Questionnaire (IPAQ; Craig et al., 2003). All assessments were performed by a trained exercise therapist and with patients in stable on medication without presence of motor-fluctuations.

In case of an allocation to the EX condition, the participants additionally underwent cardiologic screening at least 48 hours prior to the main experiment. The procedure included a graded exercise test (GXT) performed on a stationary cycle ergometer and was supervised by a cardiologist. Briefly, participants started at 25 Watts with a constant pedal rate of ≥ 60 revolutions per minute (rpm). Subsequently, load was stepwise increased by 25 Watts every 3 minutes until exhaustion, while heart rate (HR) response was recorded continuously.

### 2.4 Motor Learning Task

We used a dynamic balancing task to examine participants’ motor learning performance since balance training (i.e. postural motor learning) is a core component of motor rehabilitation in PD. (Peterson, Dijkstra, & Horak, 2016). The stabilometer is widely used to study motor learning (Lewthwaite & Wulf, 2010; Mégrot & Bardy, 2006; Wulf & Lewthwaite, 2009; Wulf, Weigelt, Poulter, & McNevin, 2003), including PD populations (Chiviacowsky, Wulf, Lewthwaite, & Campos, 2012; Sehm et al., 2014; Steib et al., 2018). Furthermore, the device has already been reported to be appropriate as an ecologically valid device to investigate learning in complex motor skills (Wulf & Shea, 2002). The device consists of a 107 x 65 cm wooden platform (stability platform; Lafayette Instrument Europe; Loughborough, United Kingdom), which is mounted on a fulcrum and has a maximum deviation of 15° to either side. Participants are required to stand with both feet on the platform and try to keep the platform as close to the horizontal as possible during a 30-second trial (Chiviacowsky, Wulf, & Wally, 2010; Steib et al., 2018; Zech et al., 2018). A millisecond timer measured time in balance, which is the time participants were able to keep the platform within ± 5° from horizontal during each 30-second trial. This common definition of time in balance was used according to current literature showing the appropriateness of this method to assess motor learning in healthy and PD populations (Chiviacowsky et al., 2010; Davlin-Pater, 2008; Steib et al., 2018; Wanner, Müller, Cristini, Pfeifer, & Steib, 2020). Since the way this motor task is instructed may affect learning outcome (Wulf & Lewthwaite, 2009), standardized formulations were used to present the task to the participants, excluding any forms of motivation or the direction of attentional focus. We further provided rigidly standardized feedback on the achieved time in balance (knowledge of results) immediately after each trial but did not give any additional information on the participants’ movement strategy. Participants were secured with a safety harness and instructed to stand in a comfortable individually chosen position (foot position was marked to ensure a standardized positioning during all tests).

On acquisition session, participants performed a familiarization trial followed by 15 practice trials (30 seconds), clustered into five blocks of three trials, with 60 seconds rest between trials and 120 seconds rest between blocks. The 1-day retention test included a warm-up trial as well as another block of three trials, whereas the 7-day retention test followed the procedure of the acquisition session (i.e. warm-up trial followed by 15 trials, clustered into 5 blocks of 3 trials). We applied only one block during the 1-day retention test in order to assess retention performance while avoiding a new learning stimulus. In contrast, during 7-day retention test, we additionally intended to assess potential performance improvements with continued practice (i.e. potential ceiling effects).

After each motor practice session (i.e. acquisition, 1-day and 7-day retention), participants completed the National Aeronautics and Space Administration-Task Load Index (NASA-TLX) (Hart & Staveland, 1988). The NASA-TLX is a visual analog scale to assess the subjective workload of a task. It consists of the six dimensions mental demand, physical demand, temporal demand, performance, effort, and frustration. Each dimension can be rated on a twenty-step bipolar scale ranging from 0 (not demanding) to 20 (extremely demanding). Moreover, the average of the six dimensions can be calculated to determine the Raw Task Load Index (RTLX), which is a measure of the overall subjective workload (Hart, 2006). The NASA-TLX has already been shown to be appropriate to assess task difficulty in balance tasks (Akizuki & Ohashi, 2015). Additionally, daytime sleepiness and sleep quality were assessed using the Epworth Sleepiness Scale (ESS; Johns, 1991) and the Pittsburgh Sleep Quality Index (PSQI; Monk et al., 1994), respectively. These tests were conducted to account for potential differences in sleep that might modulate the effects (Roig et al., 2016).

### 2.5 Exercise Protocol

During the acquisition session, participants in the EX condition performed a single exercise bout immediately (≤ 5 minutes) following motor skill practice. Briefly, the bout comprised a 5-minute warm-up (load was progressively increased until target intensity), followed by 25 minutes of pedaling at ≥ 60 rpm and an intensity of 60% maximal power output (W_max_) of the GXT. Similar protocols have been used in studies of healthy young adults, and demonstrated improved motor skill learning (Chartrand et al., 2015; Snow et al., 2016; Statton, Encarnacion, Celnik, & Bastian, 2015). Furthermore, in a recent experiment we could demonstrate that a similar exercise bout preceding motor skill practice enhanced motor memory consolidation in PD patients, indicating the appropriateness of the protocol in PD (Steib et al., 2018). HR was continuously recorded using a HR monitor (Polar Electro; Kempele; Finland). Additionally, patients’ subjective rate of perceived exertion (RPE; Borg scale 6 - 20) and blood pressure were recorded every three minutes. In case participants reported a high RPE (i.e. Borg scale ≥ 16), intensity was progressively decreased until a moderate level was reached again (i.e. Borg scale ≤ 15). According to this, load was decreased in two participants, while adjusted intensity was ≥ 50 % W_max_. In contrast, participants in the REST condition remained seated following motor skill practice for 30 minutes while they were allowed to read.

### 2.6 Statistical Analysis

All statistical analyses were performed with IBM SPSS Statistics version 26.0 and the alpha level set at p < .05. Normality, variance homogeneity and sphericity of the data were checked where appropriate.

#### 2.6.1 Participant characteristics and self-perceived task load

We compared participant characteristics between the two experimental conditions (i.e. EX vs. REST) using independent samples t-tests in normally (i.e. demographics, MDS-UPDRS-III, MoCa, IPAQ, and ESS) or Mann-Whitney-U-test in non-normally distributed (i.e. Hoehn & Yahr stage and PSQI) data.

Similarly, self-perceived task load assessed by the NASA-TLX was compared by independent samples t-test or Mann-Whitney-U-test in normally and non-normally distributed data, respectively.

#### 2.6.2 Baseline performance and memory encoding (online learning)

To ensure that both groups had the same performance at baseline, we compared the mean of the first acquisition block. Since Shapiro-Wilk test revealed non-normal distribution of the REST group (p = .010), this was done using Mann-Whitney-U-test.

Memory encoding (i.e. online skill learning) was assessed using a mixed ANOVA testing for the between-subject factor *Condition* (EX, REST) and the within-subject factor *Blocks*. Specifically, the ANOVA consisted of a 2 (EX, REST) x 5 (acquisition blocks 1-5) model.

#### 2.6.3 Memory consolidation (offline learning) and continued practice

Effects of post-practice exercise on motor memory consolidation (i.e. offline learning) were analyzed in a separate mixed ANOVA testing for the between-subject factor *Condition* (EX, REST) and the within-subject factor *Blocks*. To ensure factorizing for differences in performance at the end of skill practice, retention test scores were normalized to performance at last acquisition block (i.e. percentage change) (Lundbye-Jensen et al., 2017; Roig et al., 2012). Consequently, effect of EX on offline learning was analyzed in a 2 (EX, REST) x 2 (1-day retention block, 7-day retention block 1) model. Furthermore, since age significantly correlated with 7-day retention test offline change scores (r = - .572, p = .017), age was entered as a covariate into the model (i.e. mixed ANCOVA). In case of a significant effect, we performed pairwise comparisons for each percentage change score in a separate one-way ANCOVA accounting for age.

Continued learning on 7-day retention session was tested in a separate 2 (EX, REST) x 5 (7-day retention block 1-5 as percentage change from last acquisition block) model with performance entered as percentage change scores from last acquisition block and age as covariate (i.e. mixed ANCOVA).

## 3 Results

Data from one participant in the EX group revealed a baseline performance of > 2 SDs above the mean, and therefore was excluded from analysis. After removing the participant, the EX group consisted of eight and the REST group of nine PD patients for final analysis.

### 3.1 Participant characteristics and self-perceived task load

Participant characteristics and exercise data of the finally included PD patients are presented in table 1. The groups were not statistically different in demographics (i.e. age, height, and weight), disease severity (i.e. Hoehn & Yahr status and MDS-UPDRS-III), cognitive status (i.e. MoCa), self-reported physical activity (i.e. IPAQ), and sleep behavior (i.e. ESS and PSQI). In average, the participants in the EX group worked at 95.1 Watt (range: 60.0 - 120.0) with a heart rate of 125.6 bpm (range: 86.5 - 159.6) and a RPE of 13.6 (range: 13.0 - 14.7) during the 25-minute main part of the exercise bout.

**Table 1.** Participants’ characteristics and exercise parameters; mean ± standard deviation.

| Group | Demographics |  |  |  |  |  |  |  |  |  |  | Graded exercise test |  |  | Exercise bout<br>(25-min main part) |  |  |
| --- | --- | --- | --- | --- | --- | --- | --- | --- | --- | --- | --- | --- | --- | --- | --- | --- | --- |
|  | Gender<br>(female/<br>male) | Age<br>(yrs) | Weight<br>(kg) | Height<br>(m) | BMI | IPAQ<br>(MET-min/<br>week) | H&Y<br>stage | MDS<br>UPDRS-III | MoCA | ESS | PSQI | Watt<sub>max</sub><br>(W) | Watt<sub>max</sub><br>(W/kg) | HR<sub>max</sub><br>(bpm) | Power<br>output<br>(W) | HR<br>(bpm) | RPE |
| EX | 3/5 | 59.3<br>± 7.4 | 73.3<br>± 14.1 | 1.74<br>± 0.10 | 24.2<br>± 3.5 | 3605.1<br>± 1521.3 | 2.0<br>± 0.5 | 21.0<br>± 9.7 | 25.9<br>± 2.5 | 6.6<br>± 4.2 | 6.1<br>± 3.1 | 162.5<br>± 32.7 | 2.5<br>± 0.5 | 145.5<br>± 21.7 | 95.1<br>± 21.2 | 125.5<br>± 21.5 | 13.6<br>± 0.6 |
| REST | 4/5 | 60.8<br>± 8.8 | 84.1<br>± 13.8 | 1.74<br>± 0.05 | 27.7<br>± 4.8 | 2401.2<br>± 1599.8 | 2.0<br>± 0.3 | 21.9<br>± 5.4 | 26.2<br>± 1.9 | 8.7<br>± 4.4 | 5.9<br>± 2.6 | n/a | n/a | n/a | n/a | n/a | n/a |
| *p-value* | *n/a* | *.705* | *.131* | *.862* | *.110* | *.134* | *.743* | *.816* | *.750* | *.349* | *.815* | *n/a* | *n/a* | *n/a* | *n/a* | *n/a* | *n/a* |

EX = exercise; BMI = body mass index; IPAQ = International Physical Activity Questionnaire; H&Y = Hoehn & Yahr; MDS-UPDRS III = Unified Parkinson's Disease Rating Scale motor examination; MoCa = Montreal Cognitive Assessment; ESS = Epworth Sleepiness Scale; PSQI = Pittsburgh Sleep Quality Index; W = Watt; HR = heart rate; bpm = beats per minute; RPE = rate of perceived exertion.

Analysis of the NASA-TLX revealed no significant differences in sum scores (acquisition: T15 = .233, p = .819; 1-day retention: T15 = 1.238, p = .235; 7-day retention: T15 = .778, p = .449). Similarly, none of the sub-categories demonstrated significant group differences expect for performance during 1-day retention test with the EX group reporting a lower self-perceived performance (T15 = 2.171, p = .046) (Table 2).

**Table 2.** Self-perceived task demands.

|  | EX |  | REST |  | *p-value* |
| --- | --- | --- | --- | --- | --- |
|  | Mean | SD | Mean | SD |  |
| **Acquisition** |  |  |  |  |  |
| Mental demand | 11.9 | 4.3 | 10.6 | 6.3 | *.888* |
| Physical demand | 10.1 | 5.3 | 10.1 | 4.4 | *.995* |
| Temporal demand | 3.5 | 2.5 | 4.6 | 4.3 | *.888* |
| Performance | 10.1 | 4.4 | 9.8 | 5.6 | *.890* |
| Effort | 11.6 | 6.1 | 9.6 | 5.2 | *.462* |
| Frustration | 3.1 | 2.2 | 3.8 | 4.7 | *.673* |
| *RTLX* | 8.4 | 2.4 | 8.1 | 3.4 | *.819* |
| **Retention 1-day** |  |  |  |  |  |
| Mental demand | 9.0 | 3.9 | 8.3 | 5.6 | *.781* |
| Physical demand | 7.5 | 3.5 | 8.2 | 4.4 | *.715* |
| Temporal demand | 2.9 | 2.1 | 3.8 | 2.8 | *.466* |
| Performance | 12.6 | 3.0 | 8.8 | 4.1 | *.046\** |
| Effort | 9.5 | 4.2 | 8.1 | 4.3 | *.513* |
| Frustration | 8.1 | 4.4 | 3.8 | 4.2 | *.055* |
| *RTLX* | 8.3 | 2.1 | 6.8 | 2.7 | *.235* |
| **Retention 7-day** |  |  |  |  |  |
| Mental demand | 10.0 | 5.1 | 8.9 | 5.5 | *.674* |
| Physical demand | 12.1 | 4.6 | 8.3 | 5.3 | *.236* |
| Temporal demand | 4.1 | 2.4 | 5.9 | 3.8 | *.272* |
| Performance | 8.4 | 4.7 | 8.3 | 5.3 | *.987* |
| Effort | 11.3 | 4.7 | 9.1 | 5.9 | *.424* |
| Frustration | 6.1 | 4.5 | 4.3 | 3.2 | *.356* |
| *RTLX* | 8.7 | 2.9 | 7.5 | 3.3 | *.449* |

\* statistically significant differences; $p < .05$ ; SD = standard deviation; EX = exercise; RTLX = Raw Task Load Index.

### 3.2 Baseline performance and memory encoding (online learning)

Time in balance at first acquisition block did not significantly differ between groups (U = 27.000, p = .423) indicating similar baseline performance. ANOVA for online gains revealed a significant main effect for *Block* (F4,60 = 10.916, p < .001, η^2^*p* = .421) while no significant effect for *Condition* (F1,15 = .372, p = .551, η^2^*p* = .024) and *Condition x Block* interaction effect (F4,60 = .316, p = .866, η^2^*p* = .021), confirming task learning with a similar rate in both groups (figure 3A).

### 3.3 Memory consolidation (offline learning) and continued practice

Mixed ANCOVA for offline gains revealed no significant main effect for *Block* (F_1,14_ = 2.632, p = .127, η^2^*_p_* = .158) or *Condition* (F_1,14_ = .802, p = .386, η^2^*_p_* = .054). However, a significant *Condition x Block* interaction effect (F_1,14_ = 5.392, p = .036, η^2^*_p_* = .278) indicated that offline learning differed between groups. Subsequent separate ANCOVA for 1-day offline change scores revealed no significant difference (F_1,14_ = .650, p = .434, η^2^*_p_* = .044, EX = −8.3% vs. REST = −2.8%, figure 3B). In contrast, a significant ANCOVA for change scores to 7-day retention test suggested better offline learning in the EX group (F_1,14_ = 5.602, p = .033, η^2^*_p_* = .286, EX = 12.4% vs. REST = −5.1%, figure 3C).

We additionally analyzed skill improvement with continued practiced during 7-day retention session in a separate ANCOVA. The absence of a statistically significant main *Block* (F_4,56_ = .750, p = .562, η^2^*_p_* = .051), *Condition* (F_1,14_ = 1.318, p = .270, η^2^*_p_* = .086), or *Condition* x *Block* interaction (F_4,56_ = 1.539, p = .203, η^2^*_p_* = .099) effect indicated that participants did not improve performance with continued practice irrespective of group.

**Figure 2.**
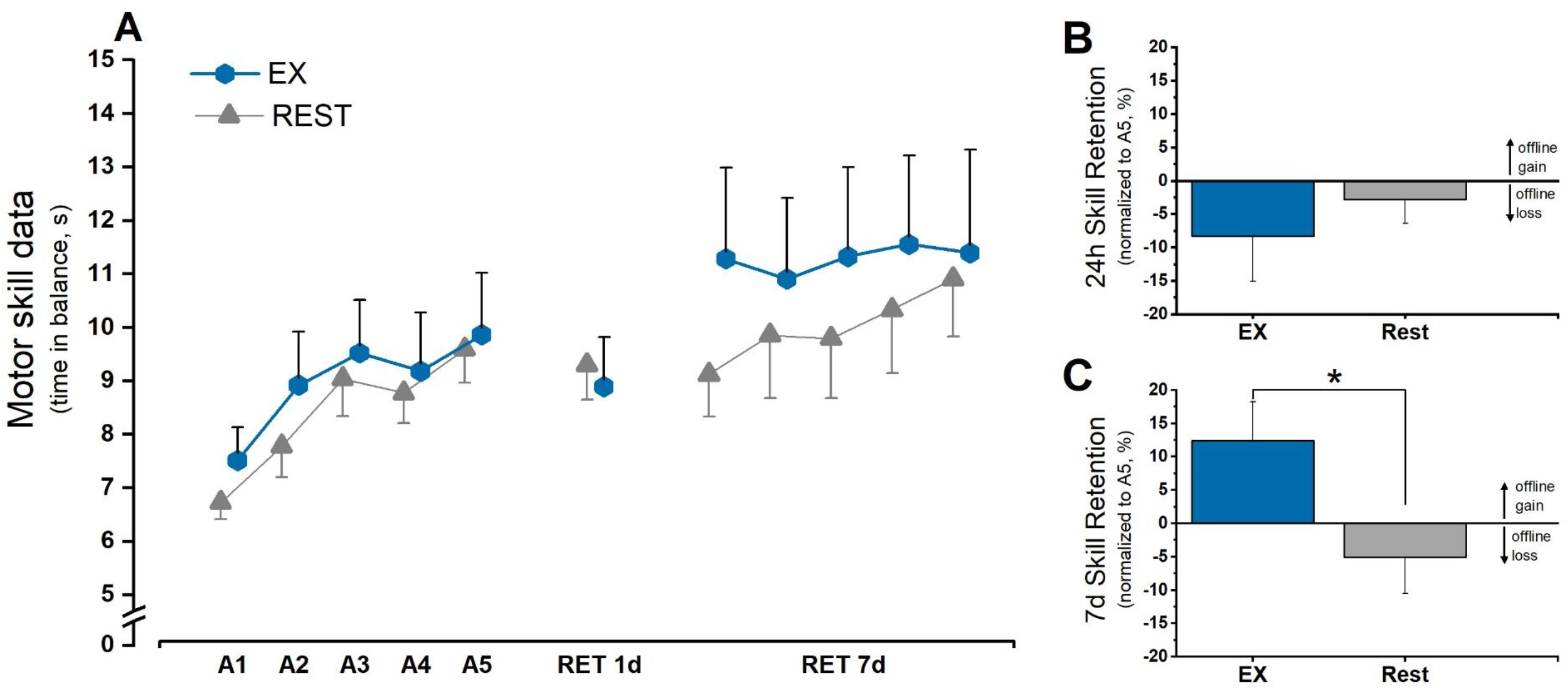
Motor skill data. **(A)** Mean motor skill performance (time in balance) during acquisition and retention (clustered in blocks of three trials); **(B)** offline skill learning illustrated as mean percentage change from last acquisition block to 1-day retention block; **(C)** offline skill learning illustrated as mean percentage change from last acquisition block to first 7-day retention block; * indicates p < .05; error bars indicate 1 standard error of the mean.

## 4 Discussion

In the present study, we aimed at investigating the effects of moderate intense cardiovascular exercise performed immediately after practicing a balance task on motor memory consolidation in PD patients. In line with our hypothesis, groups showed similar improvements during skill practice, while exercise facilitated motor memory consolidation as indicated by improved offline learning gains. Specifically, this was evident at 7-day, but not 1-day retention test.

In general, our findings are consistent with the majority of experiments in healthy individuals reporting improved offline skill learning at 7-day retention test with post-practice exercise (Ferrer-Uris, Busquets, & Angulo-Barroso, 2018; Lundbye-Jensen et al., 2017; Roig et al., 2012; Thomas, Johnsen et al., 2016; Tomporowski & Pendleton, 2018). The enhanced motor memory consolidation in PD support the promoting effects of exercise particularly on dopamine neurotransmission (Christiansen et al., 2019; Mang et al., 2017). First results from animal models as well as humans revealed increased dopamine expression and receptor binding following chronic exercise also in PD (Fisher et al., 2013; Jakowec, Wang, Holschneider, Beeler, & Petzinger, 2016; Petzinger et al., 2015; Sacheli et al., 2019). Remarkably, restoring dopamine has shown to recover neural plasticity in PD, and thus motor learning performance is suggested to be improved during peak dopamine levels (Beeler et al., 2010; Beeler, 2011; Beeler et al., 2012; Beeler, Petzinger, & Jakowec, 2013). Therefore, our results suggest an exercise-induced upregulation of dopamine as well as an improved receptor binding during critical stages of memory formation that may account for enhanced long-term memory consolidation. This finding also has important implications for motor rehabilitation purposes, since our results suggest that even acute exercise may have beneficial effects on dopamine neurotransmission. In line with this, a first study by Kelly and colleagues (2017) observed an increased substantia nigra activity following a single exercise bout in PD patients, reinforcing that already acute exercise has promoting effects on the dopaminergic system.

In addition to dopamine, an exercise-induced upregulation of neurotrophic factors, such as BDNF, has been linked to improved memory formation (Dinoff et al., 2017; Knaepen, Goekint, Heyman E.M., & Meeusen, 2010; Szuhany, Bugatti, & Otto, 2015). Studies on exercise-induced BDNF response in PD as well as other neurological diseases revealed a positive relationship (Briken et al., 2016; Gold et al., 2003; Hirsch, van Wegen, Newman, & Heyn, 2018; Morais et al., 2018). Post-practice exercise theoretically should have elevated BDNF levels during the process of memory consolidation, which might account for enhanced memory consolidation (Skriver et al., 2014). However, the exact role of exercise-induced BDNF in enhanced memory formation needs further investigation (Baird et al., 2018; Charalambous et al., 2018; Helm et al., 2017; Mang, Snow, Campbell, Ross, Colin J D, & Boyd, 2014).

Furthermore, dopamine and BDNF might indirectly mediate changes on the corticospinal level (i.e. system level of brain organization) and thereby facilitating motor memory consolidation (El-Sayes et al., 2018; Mellow et al., 2019; Singh & Staines, 2015). Findings from healthy adults showed that improved offline learning was directly linked to an exercise-induced increase in corticospinal excitability (Ostadan et al., 2016), reduction in short-interval intracortical inhibition (SICI; i.e. synaptic GABA_A_ inhibition) (Stavrinos & Coxon, 2017), and improvement of functional networks in sensorimotor areas (Dal Maso et al., 2018). Interestingly, similar changes on the system level in response to acute exercise has already been demonstrated in other neurological conditions (Chaves, Devasahayam, Kelly, Pretty, & Ploughman, 2020; Nepveu et al., 2017).

Our findings may have important implications for neurorehabilitation in PD as well as other populations with learning impairments. As aforementioned, the depletion of dopamine within the caudal basal ganglia in PD has shown to affect corticostriatal as well as thalamostriatal circuits, which are major circuits in motor learning (Petzinger et al., 2013). Specifically, these circuits are proposed to be involved in early consolidation (associative striatum) and the development of automaticity (sensorimotor striatum) (Beeler et al., 2013; Doyon et al., 2009). Consequently, learning deficits in PD have particularly seen by an impaired memory consolidation compared to age-matched nondisabled individuals (Marinelli et al., 2017). However, an elementary component of motor rehabilitation in PD is the practice-related improvement and (re-)learning of motor skills (i.e. motor learning) (Abbruzzese et al., 2016). While a first study demonstrated enhanced motor skill learning with chronic exercise in PD (Duchesne et al., 2015; Duchesne et al., 2016), our results indicate that already a single exercise bout during early memory consolidation is a promising method to counteract learning deficits. In line with this, a single post-practice exercise bout has shown to improve memory consolidation of motor as well as cognitive tasks in other populations with learning impairments, such as stroke and amnestic mild cognitive impairment (Nepveu et al., 2017; Segal, Cotman, & Cahill, 2012).

In a previous experiment, we already examined the effects of acute exercise on motor memory consolidation in PD (Steib et al., 2018). In contrast to the present study, patients performed the exercise bout immediately prior to motor skill practice. In general, our finding of an enhanced motor memory consolidation with exercise is consistent across studies. Interestingly, with exercise preceding skill practice we supposedly observed improved offline learning at 1-day retention, whereas this was not the case in the present study. This might suggest that the effects critically depend on exercise timing (Roig et al., 2016). However, participants in the pre-practice exercise condition demonstrated diminished skill improvements during late practice (i.e. block 4-5). Based on the present findings (i.e. no enhanced 1-day retention performance) it cannot be ruled out that the improved offline learning in our previous study might reflect fatigue during the end of skill practice (i.e. diminished performance), which is reduced during 1-day retention test (Robertson, 2019).

We found effects of exercise on 7-day, but not on 1-day retention. This partly corresponds to studies on healthy individuals showing stronger effects of post-practice exercise on long-term retention performance (Lundbye-Jensen et al., 2017; Thomas, Johnsen et al., 2016; Tomporowski & Pendleton, 2018). It has been proposed that the consolidation of a newly acquired skill requires time (Dudai, 2012). Accordingly, the promoting effects of exercise might particularly become visible after a longer latency period since plasticity-promoting effects of exercise have more time to stabilize the motor engram.

Interestingly, Nepveu and colleagues (2017) recently found improved 1-day retention performance of a fine-motor task (i.e. visuomotor tracking) with post-practice exercise in stroke survivors. The discrepancy to our results might be attributed to the different motor tasks used (Roig et al., 2016). In accordance to the NASA-TLX, the physical demands of our task were relatively high, indicating higher loads on the motor system (table 2). It has already been suggested that exercise might particularly facilitate the cognitive components of motor performance, and effects are less visible in highly motor challenging tasks (Baird et al., 2018). In line with this interpretation, Charalambous and colleagues (2018) could not observe enhanced 1-day memory consolidation of gait adaptation to split-belt treadmill walking with exercise in stroke survivors. This again questions the generalizability of findings from simple motor skills and reinforces the need for studying more complex skills (e.g., balance, gait) in order to draw ecologically valid conclusions for (neuro-)rehabilitation purposes. Alternatively, negative consequences of exercise-induced physical fatigue might still be existent during 1-day retention test, negatively affecting retention performance (Roig et al., 2016). These detrimental effects of fatigue may particularly be present in clinical populations (e.g., PD patients), as our participants were of higher age and show motor impairments. Furthermore, using an exercise type (i.e. cycling) involving muscle groups similar to the motor learning task (i.e. balancing) might even increase fatigue effects. Remarkably, the existence of fatigue during 1-day retention session would be supported by the significantly lower self-perceived task performance in the EX group (table 2). The performance item provides information about the self-perceived quality of movement and has shown to be the best predictor for task difficulty of balance tasks (Akizuki & Ohashi, 2015). Hence, the lower self-rated performance in the EX group may indicate higher task difficulty as consequence of fatigue.

## 5 Limitations

We could not collect neurophysiological correlates, which would have allowed us to directly relate our results to exercise-induced neuroplastic processes. Further, the lack of improvements in performance with continued practice may indicate ceiling effects on 7-day retention test. Even though not statistically significant, visual observation suggests that missing improvements were particularly present in the EX group. Consequently, potential ceiling effects may impeded larger offline gains in the EX group.

## 6 Conclusion

Results of the present pilot study support that cardiovascular exercise is effective in facilitating memory consolidation in individuals with motor and cognitive deficits. Specifically, our results suggest that even a single moderate intense bout performed immediately following skill practice improves long-term motor memory consolidation in PD patients. This finding may point at exercise-induced beneficial effects on the dopaminergic system. Furthermore, our results have promising implications for neurorehabilitation, since motor learning processes are a core component in motor recovery, but learning ability is often impaired. Exercise appears to optimize the memory formation processes, and thus counteract learning deficits. Future work should focus on the acute effects of exercise on dopamine neurotransmission, investigate optimal exercise parameters (e.g., intensity and duration), and try to transfer these promising effects to other populations with learning dysfunctions.

## Acknowledgements

We would like to thank Dr. Harald Erxleben from the Neurologische Klinik at Sana-Krankenhaus Rummelsberg/Nuremberg for his assistance in participant recruitment as well as Dr. Leonard Frauenberger from the Division of Sport and Exercise Medicine at the Department of Sport Science and Sport of the FAU and Dr. Mohammed Asker Hasan from the Herzkatheterlabor at Sana-Krankenhaus Rummelsberg/Nuremberg for their medical advice and support in cardiac screening and exercise stress testing.

